# Pavement cells and the topology puzzle

**DOI:** 10.1101/160762

**Authors:** Ross Carter, Yara E. Sánchez-Corrales, Verônica A. Grieneisen, Athanasius F. M. Marée

## Abstract

D’Arcy Thompson emphasised the importance of surface tension as a potential driving force in establishing cell shape and topology within tissues. Leaf epidermal pavement cells grow into jigsaw-piece shapes, highly deviating from such classical forms. We investigate the topology of developing *Arabidopsis* leaves composed solely of pavement cells. Image analysis of around 50,000 cells reveals a clear and unique topological signature, deviating from previously studied epidermal tissues. This topological distribution is however established early during leaf development, already before the typical pavement cell shapes emerge, with topological homestasis maintained throughout growth and unaltered between division and maturation zones. Simulating graph models, we identify a heuristic cellular division rule that reproduces the observed topology. Our parsimonious model predicts how and when cells effectively place their division plane with respect to their neighbours. We verify the predicted dynamics through in vivo tracking of 800 mitotic events, and conclude that the distinct topology is not a direct consequence of the jigsaw-like shape of the cells, but rather owes itself to a strongly life-history-driven process, with limited impact from cell surface mechanics.

**Summary statement**

Development of the *Arabidopsis* leaf epidermis topology is driven by deceptively simple rules of cell division, independent of surface tension, cell size and, often complex, cell shape.

## Introduction

Spatio-temporal control of cell growth and division is involved in the generation of tissue shape during development. Tissue shape is likewise affected by biophysical interactions between individual cells within the tessellated context that also modify the typical lengths between cellular contacts, the curvature between cells, and cellular arrangements. Two-dimensional layers of cells offer an ideal model system to investigate the cross-scale processes involved, as extensively studied by D’ Arcy Thompson (Thompson, 1917). One way to do so is by analysing the cellular topology. In “ On Growth and Form” D’ Arcy Thompson explains how cellular division rules and surface-tension minimising processes acting upon cells within tissues should yield characteristic topologies, i.e., specific distributions regarding the number of neighbouring cells, which he points out can be regarded as fingerprints of the underlying forces guiding cellular behaviour. Most of his examples refer to biological tissues that resemble foam, with geometries that are strikingly honey-comb like. An example of such a tissue is the *Drosophila* epidermis shown in Figure 1A. It is widely accepted that in cellular materials in which superficial tension dominates, cells tend to acquire a hexagonal shape, obtaining an average of six neighbours. Such pattern generation can also be observed in artificial tissue (Figure 1B, Farhadifar et al., 2007; Lecuit and Lenne, 2007; Lewis, 1931; Magno et al., 2015; Thompson, 1917). This topology minimises surface area for a given number of cells and optimises the packing density (Durand, 2015; Weaire and Rivier, 1984). But D’ Arcy Thompson also drew attention to a few “ misfits” in the zoo of cell and tissue shapes, such as the endothelium of blood-vessels (Figure 1C, a), the epithelial cells that form the gills of a mussel, and, finally, the epidermal pavement cells of plant leaves (Figure 1C, b-c, D). The odd sinusoidal features which such cells present seem to defy the principles of surface minimisation. Nevertheless, D’ Arcy Thompson offers an explanation by means of analogy: “ If we make a froth of white-of-egg upon a stretched sheet of rubber, the cells of the froth will tend to assume their normal hexagonal pattern; but relax the elastic membrane, and the cell-walls are thrown into beautiful sinuous or wavy folds” (Thompson, 1917). He argues that such buckling forces could be operating in the animal epithelium, and could account for the sinusoidal cellular interfaces. Yet, for the jigsaw-like shape of plant pavement cells, he only briefly mentions that “ the more coarsely sinuous outlines of the epithelium in many plants is another story, and not so easily accounted for” (Thompson, 1917).

**Figure 1:**
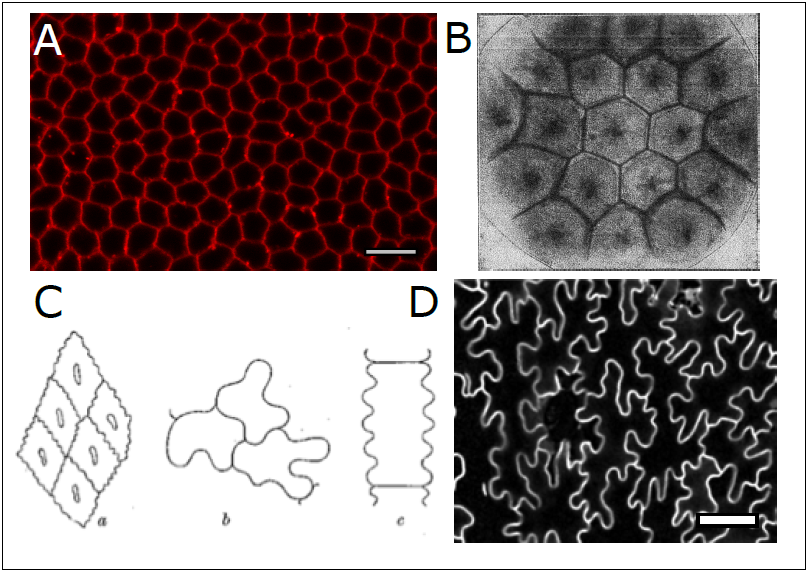
Foam-like cells and puzzle-like cells. Biological tissues, such as *Drosophila* epithelium (A), can adopt geometric resemblance to non-biological materials such as ‘ artificial tissue’ in which surface tension processes dominate (B), here formed by coloured drops of a solution diffusing on a less dense solution of a same salt (Thompson, 1917). (C) Cells presenting sinuous outlines, as illustrated in Thompson (1917), Fig. 186: endothelium of a bloodvessel (a); and plant tissues *Impatiens* (b) and *Festuca* (c). (D) Confocal image of the PCs in mature *Arabidopsis* leaves, which have grown into jigsaw-puzzle-like shapes.

Recent molecular and biophysical studies have indeed shown that pavement cell (PC) shapes arise due to active internal processes driving anisotropic growth, as a consequence of intracellular patterning within the individual cells (Fu et al., 2005, 2009; Gu et al., 2006). The internal patterning involves feedbacks between Rhos of Plants and cytoskeletal elements (Fu et al., 2005, 2009; Grieneisen, 2009; Grieneisen et al., 2013), causing modifications in the structural properties of the cell walls, thereby triggering lobe and indentation formation between those cells (Fu et al., 2009).

Cellular topology arises from the interplay between the way cell division is organised and the biophysical interactions between the neighbouring cells. Cell divisions modify the topological constitution, while biophysical interactions between cells can change the topology either directly, by triggering neighbourhood changes between cells, or indirectly, through modifications of cell interface length or cell shape, in its turn affecting, in topological terms, the next cell division plane. While for plant tissues neighbourhood changes are considered to be impossible, the indirect effect of the biophysical interactions can play an important role. In this context, the resultant shapes of pavement cells seem to be little influenced by the surface-tension driven processes that characterise tissues with hexagonal symmetries, instead originating in a complex biologically-controlled manner. Given that the surface-tension driven processes generate a very clear topological fingerprint, we asked if tissues that are only composed of pavement cells would likewise display a unique topological composition, distinct to the ones observed for the more “ honey-comb” - like patterns. It would thus allow to determine the relative contribution and importance of cell-surface-mechanics-driven mechanisms vs life-history-driven mechanisms in the establishment of the tissue topology, which, in its turn, constraints the potential diversity in cell shapes an sizes.

## Results

The *Arabidopsis* leaf epidermis provides an ideal system to observe neighbourhood topology within a developing tissue, due to the fact that its leaves are relatively flat during the development and the epithelium is composed out of a singe layer of very thin (quasi-2D) cells. Since we here wish to focus on the tessellations that arise amongst pavement cells (PCs), we specifically grew Arabidopsis *speechless* (*spch*) mutant lines, in which stomatal lineages are not present (for details see Materials and Methods). These lines were crossed with a membrane marker, to allow us to visualise the boundaries between cells using confocal microscopy (see Materials and Methods). We followed the growth of several leaves over extended periods, one of them consecutively for a period of nearly 16 days.

### Spatial topological patterns over the leaf

As eluded to, topology and geometry are two important aspects to consider when unravelling the mechanisms guiding the development of epithelia. Geometry refers to the shape and size of the cells that make up the epithelium, whereas topology refers to the connectivity of the cells within the tissue, i.e., the number of neighbours of each cell.

The geometry of pavement cells at early stages of leaf development is fairly isotropic (Figure 2A, B), with cells along the midline of the leaf being more anisotropically shaped. As developmental time progresses, cells grow and develop into the unique “ jigsaw-puzzle” - like shapes, in a graded fashion from the tip of the leaf to the direction of the base (Figure 2C, D). We first analysed if the development of such characteristic shapes also affects the topology within the tissue, either when these cells first arise within the young leaf, or later on during the progressive development of the leaf. To screen for possible topological patterns at different developmental stages, as well as over the tissue itself, we colour-coded the segmented cells of the epidermis, indicating for each cell its number of neighbours. We here show the results for two distinct time points, one early (time-point 0, at 175.17 h after stratification) and one later (time-point 9, 286.50 h after stratification) during the leaf development (Figure 2B, D). When cells at the boundary or with an incomplete set of segmented neighbours are excluded, as is done here, then a perfectly homogeneous and honey-combed shaped cellular material would appear “ white”, indicating that each cell would have six neighbours, see colour bars in Figure 2. Instead, we find that topology differs from cell to cell. Cells with less than six neighbours are shown through a brown colour spectrum, while those with more than six neighbours are depicted using increasing shades of green. The images confer for both developmental times a preference for a general (but not strict) neighbourhood number of six. Both brownish and greenish colours are present to a high degree in an intermixed fashion in these plots, indicating the broad distribution in cell topologies that exist over the leaf. Moreover, there seems not to be any noticeable spatial structuring in the topology over the tissue itself, except for a clustering of larger neighbourhood numbers along the mid vein at the later stage, coinciding with an elongation of the cells that compose the tissue in that region.

**Figure 2:**
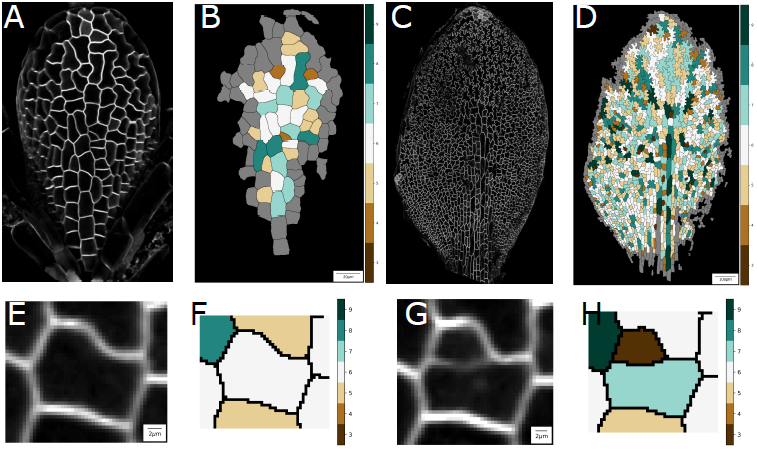
Topology across the leaf, generated through divisions. (A–D) The Arabidopsis *spch* leaf epidermis at an early time point (A, B), corresponding to time-point 0 at 175.17 h after stratification, and a later time point (C, D), corresponding to time-point 9, 286.50 h after stratification. (B, D) Heat maps of the number of neighbours of each cell, for the leaves shown in A and C, respectively. (E–H) Divisions influence local topology. A cell is shown at the time point before it divides (E), imaged at time-point 5,220.42 h after stratification, with number of neighbours indicated through a heat map (F). After division, 12.2 h later, at time point 6, the cell presents an altered neighbourhood, as indicated through the heat map (G).

### Topological distributions are conserved at different developmental stages and within different zones

Within plant systems, in which neighbourhood changes only occur in rare events (Thompson, 1917), topology is a direct consequence of the previous cell divisions (Mombach et al., 1990). The way in which a cell divides directly affects the topological constitution of its offspring, as well as the topology within its local neighbourhood (Delannay and Le Caër, 1994). This, however, does not imply that biophysics and cell surface mechanics are not involved, since the shapes cells take up after division are considered to be an important determinant for how the next cell division will be taking place. To illustrate how cell divisions affect topology, we tracked a particular cell over time, indicating how its division affects the topology of its daughter cells as well as of its neighbours (Figure 2E–H). The mother cell, which originally has six neighbours, gives rise to two daughter cells, which have seven and three neighbours, respectively. The sum of the number of neighbours of the daughter cells after a cell division is always *n* + 4, independent of the number before cell division, i.e., on average the daughter cells have 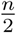 + 2 neighbours. Consequently, cells with three neighbours on average gain neighbours due to cell division, while cells with more than four neighbours on average loose neighbours due to cell division, this effect being more dramatic when the number of neighbours is high. Note that two of the neighbours of the mother cell also change topology (Figure 2E–H). For one cell the number of neighbours increases from six to seven, for the other cell it increases from five to six. Since cell division always causes a total increase of two, the neighbouring cells will, on average, have their number of neighbours increased by 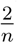.

Given that cell divisions during leaf development dynamically change in a temporally and spatially controlled manner (Andriankaja et al., 2012; Asl et al., 2011; Kazama et al., 2010), we next investigated how cell topology distributions alter during leaf development. We therefore analysed in detail the topological distribution of a young leaf that is still large enough to be composed out of a sufficiently large number of cells to allow for meaningful distributions to be made, namely at time-point 2, 193.25 h after stratification. We compared its topological distribution with that of a leaf at a more advanced developmental stage, namely at time-point 9, 286.50 h after stratification. The spatial distributions in neighbourhood numbers across the tissue (Figure 3A, B) again do not reveal any noticeable patterning, with the exception of a consistent tendency for higher neighbourhood numbers to appear close to the mid vein, with the cells in this region clearly being more elongated. Surprisingly, the topological distribution for the entire cell population is unaltered at these different time points, bearing a characteristic profile (Figure 3C). It therefore seems that the cell population within the leaf tissue establishes a topological homoeostasis, even though the geometry of the cells changes considerably over these stages, the number of cells is still rapidly increasing, and cell proliferation dramatically varies between different parts of the leaf.

**Figure 3:**
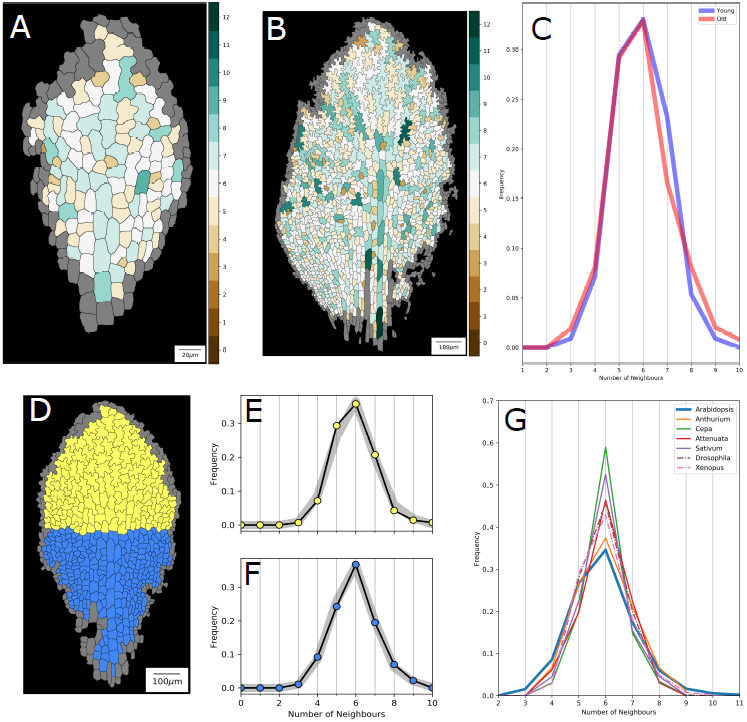
Topological distributions over time and space. (A, B) Tissue topology. A heat map of the number of neighbours for each cell for a leaf at time-point 2, 193.25 h after stratification (A), and at time-point 9, 286.50 h after stratification (B). (C) Plot comparing the neighbour frequency distributions for the ‘ young’ (A) and ‘ old’ (B) leaf. Note that the features of the distribution are conserved. (D) Meristematic (blue) and differentiation (yellow) zone of a leaf at time point 6, 232.62 h after stratification. (E) Frequency distribution for the yellow cells, which are more complex-shaped cells, compared to the distribution for an aggregate dataset consisting of 50,000 leaf cells (grey).(F) Frequency distribution for the blue, less complex and dividing cells, compared to the aggregate leaf distribution (grey). (G) Average frequency distributions for a range of animal epidermal (dotted lines, Gibson et al., 2006) and plant epidermal (solid lines, Mombach et al., 1990) tissues of a number of different species. Note that the aggregate PC dataset presents the least frequent 6-sided neighbourhood, as well as the most frequent 5- and 4-sidedness.

We therefore next analysed if we could instead distinguish between the topological distributions for the distinct cell populations within a single tissue. This was done by contrasting the topological distribution of the proximal differentiating cells to that of the dividing smaller cells at the base of the leaf (Figure 3D). This analysis is strongly linked to our understanding of how leaves develop: while cells divide at a fast rate at the base of the leaf, they stop dividing proximally, thereafter only expanding and forming complex cell shapes (Andriankaja et al., 2012). Given that the (development of the) topology is directly linked to the cell divisions, an active dividing tissue might still be presenting a different topology compared to tissue that has fully matured. Both populations, however, present very similar topological profiles, with a relatively low peak at six and broad ‘ shoulders’, including a characteristic skewness to smaller neighbourhood numbers, i.e., a high level of five neighbours and still significant fractions of four and three neighbours (Figure 3E, F). From the observation that the topological distribution is robust over developmental time and conserved between the functionally distinct leaf zones, we conclude that the total number of mitotic rounds that cells undergo does not seem to influence the topological distribution. This suggests that the tissue rapidly reaches a topological ‘ steady state’, with the topology not being affected by whether or not additional rounds of divisions take place. It also suggests that the manner in which the divisions take place should not depend on developmental time nor on the location within the leaf.

Given that the subsequent cellular development into highly complex shapes does not impact the topological distribution, we asked if these distributions therefore resemble other measured topological distributions of animal and plant tissues that do not manifest such complex shape changes (Figure 3G). Using such a cross-species comparison, based upon available published measurements (Gibson et al., 2006; Mombach et al., 1990), we can make several observations. Firstly, and perhaps not surprisingly, *Arabidopsis* pavement cells present a topological distribution which is clearly distinct from *Drosophila*. The *Drosophila* imaginal disc is a classical example of an epithelial system presenting a “ surfacetension-driven” topological signature, with a tissue of roughly equally sized and isotropic cells in which equal tensions between the cell membranes tend to relax all cells into hexagonal symmetries (Farhadifar et al., 2007; Lecuit and Lenne, 2007; Sánchez-Gutiérrez et al., 2016), as discussed extensively in Thompson (1917). In fact, all animal epidermal tissues quantified in Gibson et al. (2006); Gibson and Gibson (2009); Li et al. (2012); Sá nchez-Gutié rrez et al. (2016) present a much higher peak of six-sided cells than found for the PC tissue analysed here. But the PC tissue also distinguishes itself topologically from other plant epithelia, as reported in Mombach et al. (1990), such as *Cepa*, *Sativum* and *Attenuata* (Figure 3G). In fact, these tissues (as shown in Mombach et al., 1990) present much more ordered and staggered brick-like patterning, which again leads to much higher fractions of six-sided cells than found for our PCs. In fact, none of the plant epithelia as quantified by Korn and Spalding (1973); Lewis (1928); Mombach et al. (1990); Sahlin and Jönsson (2010) or animal epithelia as quantified by Gibson et al. (2006); Li et al. (2012); Sá nchez-Gutié rez et al. (2016) present the characteristic PC cell topologies (nor, for that matter, geometries). It is unlikely, however, that the mechanisms which drive PC shape formation drive these topological differences directly, given that the PC tissues already present their typical topological distributions prior to the jigsaw shapes arising (Figure 3A and F). As an alternative hypothesis we therefore queried if the tissue’ s unique topological distribution could be captured by a set of well-defined topological division rules instead.

### Cell division model

We chose to next adopt a purely topological test to determine to what extent the steady state topological distribution of the pavement cells in the *Arabidopsis* tissue (Figure 4A) could be accounted for by the cell division life-history only. Our representation only describes neighbourhood connections. It explicitly ignores cell shape, therefore serving as a null hypothesis that cell shape and size does not play any role. In such an approach, the epithelial tissue can be abstracted to be an undirected graph, in which cells are considered to be nodes and links to neighbours are considered to be edges. A cellular division, such as illustrated in Figure 4B, adds a cell wall and two daughter cells to the tissue, increasing the nodes and altering the edges of the graph appropriately. Intuitively it might seem nonsensical to attempt to capture the topological relationships of such an intricate system as the *Arabidopsis* PC tissue through simple division rules that are solely based on topological rules. Ignoring cell size and shape is in stark contrast with the tradition to phrase plant cell division rules in terms of their shape, such as, for example, the shortest wall algorithm, Errera’ s rule, strain-based rules, etc (Sahlin and Jönsson, 2010; Thompson, 1917). We nevertheless tested several basic topological divisions rules, performing simulations to evaluate to which extent their resulting distributions could, or could not, capture the observed robust topological distribution.

**Figure 4:**
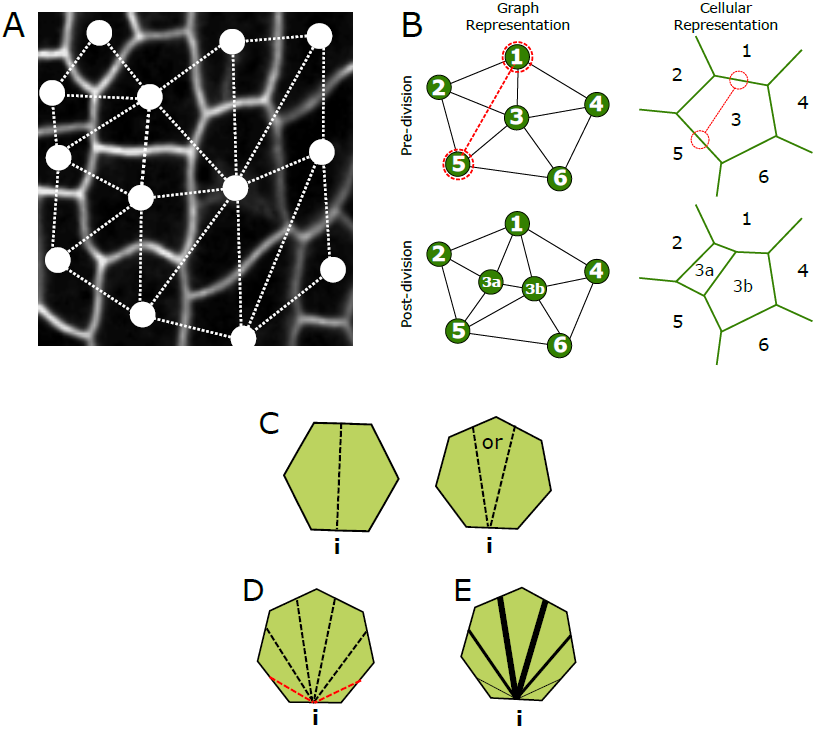
Graph model for cell divisions. (A) Network/graph representation superimposed to an actual tissue. Cells are represented by nodes, neighbours are represented by links. (B) Schematic of division implementation within the graph model (left), and its equivalent in space-embedded cellular representation (right). (C—E) The three different simulated division rules. (C) Equal split: neighbours are equally split between the two daughter cells if the cell has an even number of neighbours (*i*). Otherwise a random choice is made regarding the remaining neighbour (i´). (D) Random rule: there is an equal chance for neighbours to be split in any ratio. (E) Pascal rule: neighbour splitting follows a binomial probability, with splits in equal neighbour numbers being more favourable.

Simulations take the form of operations on a two-dimensional network. Given that four-way cell junctions are biophysically avoided and mathematically form an infinitely small subset of how junctions can be formed (Thompson, 1917), we do not allow them to be formed within the model. From this constraint, if follows that the number of neighbours of each cell in this description is equivalent to its number of sides. With such a purely topological description, all that is needed to completely define the process and consequences of cell division, is to specify which cell is dividing and the two neighbouring cells’ facets that are facing the division plane. The placement of the division plane depends upon which specific topological cell division rule is chosen. For the purpose of this study, we consider three possible ways cell division can be organised, namely an equal split division plane (Figure 4C, “ equal split”), a randomly oriented division plane (Figure 4D, “ random rule”), and a binomially weighted division plane (Figure 4E, “ pascal rule”). The first scenario, the equal split rule, implies that the new cell wall will deterministically form in such a way to equally distribute the neighbours of the mother between the daughter cells. For an even number of neighbours there is only one possibility of such a split (Figure 4C left); for an uneven number we randomly choose one of the two possibilities, with one of the two daughter cells ending up with one more neighbour than the other daughter cell (Figure 4C right). The randomly orientated division plane rule (Figure 4D) captures the situation in which there are no topological pressures whatsoever operating on the choice of the division plane location, and that any combination is equally likely. Finally, the binomially weighted division plane case (Figure 4E), the pascal rule, lies in between the other two rules: it considers it more likely that cells divide in such a manner to equally distribute neighbours between both daughters. However, in a probabilistically decreasing manner, it is still possible to asymmetrically distribute the neighbours between the cells. One way to mechanistically interpret the latter division rule, is that cells divide in two equal parts, the new cell wall connecting two different neighbouring cells, and all other neighbouring cells having an equal and independent likelihood to be adjacent to either one of the newly formed daughter cells (see also Gibson et al., 2006). This is the most likely scenario when cells divide more or less in two equal halves while the interface lengths with the neighbouring cells are randomly distributed. In all three division rules considered, we assume the orientation of division to be completely random.

Running iterative rounds of any of these three rules for each cell on the graph quickly generates steady state distributions for the final neighbour number frequencies. Surprisingly, our experimental data closely resembles that of the equal split divisions, and is qualitatively different from both the random and pascal rules (Figure 5A).

**Figure 5:**
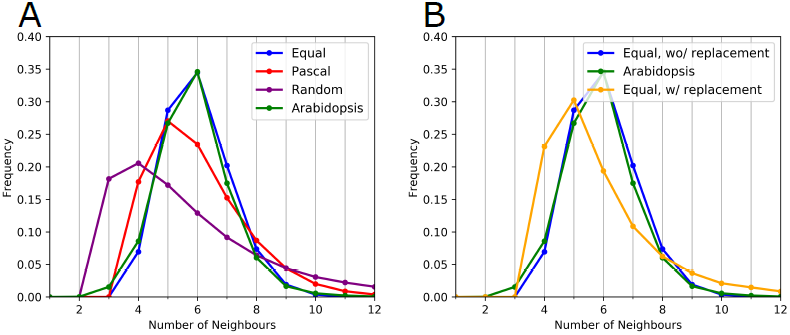
Topological distributions resulting from underlying rules. (A) Frequency distributions resulting from the different division rules compared to the aggregated *Arabidopsis* PC tissue data. Here, all rules were implemented using the ‘ without replacement’ cell division rule. The mean and the variance, μ _2_ =< *n*^^2^^ > - < *n* > ^^2^^, of these topological distributions were μ_1_ = 5.999, μ_2_ = 1.312 (Equal:blue); μ_1_ = 5.999,μ_2_ = 2.695 (Pascal:red); μ_1_ = 5.999, μ_2_ = 10.334 (Random:purple). (B) Frequency distributions for the different ‘ replacement’ division implementations, ‘ with replacement’ (yellow) and ‘ without replacement’ (blue), compared to our aggregate *Arabidopsis* data, in both cases using the Equal division rule.

Moreover, there are two main ways of implementing division rounds within the graph model simulations. One method is to consider that each cell in the tissue performs a single division during each round of divisions. In the model we coin this division ‘ without replacement’. Alternatively, the cell to divide can each time be randomly selected, its daughter cells after the division to be considered with equal probability as all other cells for the next division event. We coin such a division rule ‘ with replacement’. The latter implementation (division ‘ with replacement’) implies that some cells will undergo more divisions than the number of iteration rounds, while others cells divide less frequently, in case they get selected less often. When analysing the resultant neighbourhood frequency distributions, it becomes clear that the Equal split rule implemented ‘ without replacement’ closely matches the experimental data, while the ‘ with replacement’ version presents a distribution which is shifted to the left and broader compared to the experimental data (Figure 5B). No other combination reproduces the experimental data. These results imply that within any local region of the leaf tissue PCs should have undergone the same number of divisions and therefore be dividing with more-or-less the same frequency, i.e., the mitotic cycle within local neighbourhoods should be highly comparable. This is consistent with studies on wildtype *Arabidopsis* leaves in which average cell cycle time was found to be constant (Asl et al., 2011), and also in accordance to the manner in which division zones are structured and change over time and space (Andriankaja et al., 2012; Kazama et al., 2010). Nevertheless, it is not obvious if tight control on cell cycle and division zone would indeed lead to the same number of divisions for each cell. Not only is a certain level of random fluctuations to be expected if cells are not actively ‘ counting’ their number of divisions, but it would also imply that the large observed variation in final cell sizes cannot be explained by a certain level of spread in cell division rounds, currently the most straightforward explanation.

Gibson et al. (2006) presented a simple Markov model, entirely in terms of cell fractions, in which the divisions of cells and their neighbours causes loss and gain in the number of neighbours. They showed that a population that divides following a Pascal division rule, which moreover explicitly excludes triangular cells, will on average end up with a population of 28% pentagons, 46% hexagons, 20% heptagons and 4% higher order cells. In contrast, our leaf data shows a significant percentage of four-sided cells (7%), approximately the same number of 5-sided cells (27%), fewer 6-sided cells (39%) and around the same levels of 7- and 8-sided cells.

### In vivo tracking of underlying topological division relations

The simplicity of the topological division rule and the obvious over-simplification involved when abstracting cell division behaviour to a mere consideration regarding neighbourhood relations only, raises the question as to how to interpret the surprisingly close match between the resultant profiles. Could it be that altogether different, more complex, division rules are operating that do not obey the topological rules of the model, but still give rise to the same resultant topological distribution? To exclude such an explanation, we directly followed the *in vivo* division events themselves, by tracking 806 cell divisions and quantifying the pre- and post-mitotic neighbourhood number distributions of the mother cell and the resultant daughter cells involved. These topological changes can be captured into a matrix relating neighbourhood number probabilities of the resultant daughters cells to the original neighbourhood count of the mother cell. By comparing the matrices derived from the different division rules (Figure 6A, C, D) to the matrix derived from the *in vivo* tracking (Figure 6B), we could verify that also on the micro-rule level the ‘ Equal split’ closely and best resembles the experimental data.

**Figure 6:**
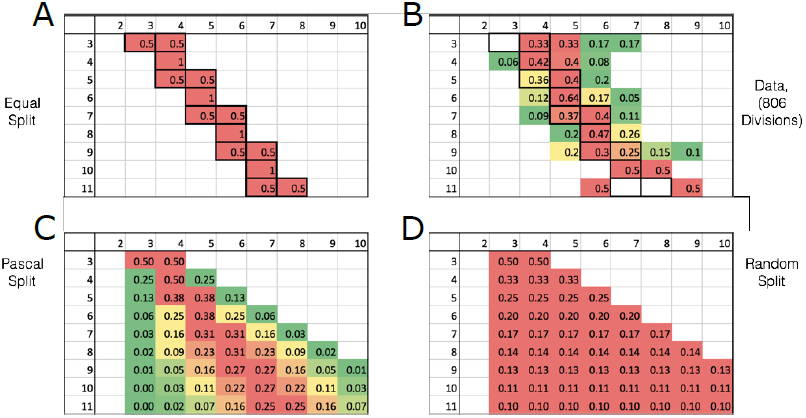
Comparing theoretical topological division rules to the experimental data. Division matrices for postmitotic neighbour number likelihood for the different division rules as well as the experimental data, with a cell’ s pre-mitotic neighbour number along the edge and the post-mitotic neighbour number of a daughter cell along the top. The table entry denotes the probability that a cell with a given number of neighbours gives rise to a daughter cell with a certain number of neighbours. As a visual guidance, green to red colour coding indicates relative likelihood for a given premitotic neighbour number. (A) Equal division rule. (B) *Arabidopsis* PC data. (C) Pascal rule. (D) Random split.

### Breaking the Law: From Aboav-Weaire back to Lewis’

The observation that the PC topology can be described well as being generated by a simple set of topological rules, combined with the observed division events themselves being very similar to the topological divisions as implemented in the model, evokes the question as to the role, if any, of cell geometry and cell shape during the developmental process. We start probing potential additional regulatory processes involved in the topological outcome by firstly comparing the topological properties of the neighbours of a given PC with a certain topology and contrasting the observed profiles to the characteristic profiles of other, non-biological cellular material. In short, we test if the Aboav-Weaire Law holds for our PC tissue. In its most basic form, Aboav empirically found (Aboav, 1970; Chiu, 1995), for a range of non-biological cellular and granular materials, that

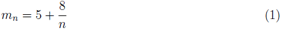

where *m*_*n*_ refers to the average number of edges of a randomly chosen cell neighbouring a cell with *n* edges (Aboav, 1970). Plotting *m*_*n*_ against *n* for our data, we observe that the data obeys the general trend that cells with higher number of neighbours tend to be surround by cells with, on average, less neighbours, in accordance with Equation 1 (Figure 7A). However, Aboav-Weaire’ s Law consistently overestimates this average as compared to our experimental data, and is also qualitatively divergent from the experimental data at *n* = 3. When analysing the same relationship for our topological models (Figure S1), we find that they also deviate from Aboav-Weaire’ s Law, which implies that, at least the set of models analysed here cannot underlie an Aboav-Weair-like relationship. The experimental data again most closely matches the Equal division rule. Nevertheless, unlike the previous results presented in this paper, we now observe a very clear discrepancy between the experimental data and the topological model, with the experimental data showing a much more pronounced relationship between neighbour number and the number of neighbour’ s neighbours. This suggests that the mechanisms underlying the second-order neighbourhood topology cannot be captured by a very basic topological rule only.

**Figure 7:**
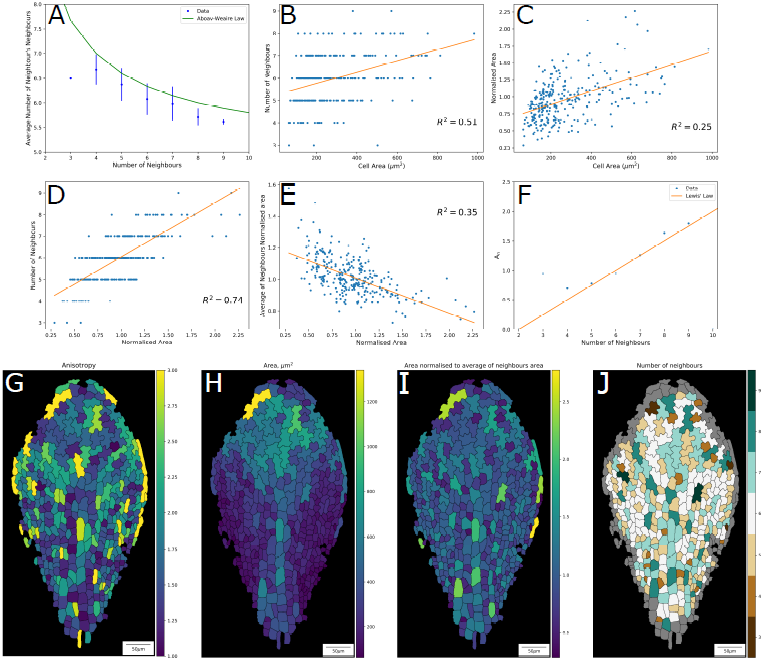
Relating topology and size. (A–F) Plots linking topological relationships and cell size for all cells of a leaf imaged at time-point 5, 220.42 h after stratification. (A) number of neighbours vs average number of neighbour’ s neighbours, with Aboav-Weaire’ s Law superimposed (green line). (B) Cell area vs number of neighbours. (C) Cell area vs normalised area (cell area/average of neighbours’ cell area). (D) Normalised area vs number of neighbours. (E) Normalised area vs average of neighbours’ normalised area (excluding the central cell itself). (F) Average area of a cells with *n* neighbours against neighbourhood number, with Lewis’ Law superimposed (orange line). (G–J) Heat maps showing cell shape properties distributed over the leaf. Heat map plots showing (G) anisotropy (major axis/minor axis); (H) cell area; (I) cell area normalised to average of neighbours; and (J) number of neighbours. Colour bar for each image shown to the right of the image. Orange lines in (B–E) indicate the linear regression line, with corresponding *R*^^2^^ value indicated within each panel.

One way to intuitively interpret the experimentally observed relationship is that the probability of cells having more neighbours is directly linked to these cells covering a larger than normal area in space, while their neighbours are covering a smaller than normal area, hence having less neighbours. Indeed, Peshkin et al. (1991) have shown that Aboav’ s law could be derived using the Maximal Entropy Principal, arguing that from a statistical perspective, larger cells tend to neighbour on average smaller ones. To verify for the experimental data the first assumption, namely that having more neighbours can indeed be linked to being larger, we next plotted the number of neighbours against cell size, but only roughly found this basic trend back (Figure 7B). The graph is very scattered (*R*^^2^^=0.51), and the more-or-less continuous variation in size over the tissue (Figure 7H) does not render the first part of the explanation robust enough to start accounting for the observed AW relationship.

Realising that the large variations in cell area over the tissue might obfuscate the relationship, we next plotted the areal ratio between each cell and that of its neighbours (which we termed ‘ normalized area’), as a function the absolute size of each cell (Figure 7C). This, however, still does not provide a strong correlation, although big cells do tend to be bigger than their neighbours, and small cells tend to be smaller than their neighbours. If, however, the number of neighbours is plotted against the normalised area (Figure 7D), a much stronger correlation emerges. In short, the number of neighbours clearly correlates with the normalised area of each cell (i.e., how much larger or smaller it is than its neighbours). This result means that within the PC tissue a cell which is relatively big in comparison to its neighbours will have, on average, more neighbours. Conversely, by being smaller than its neighbours, the cell tends to have less neighbours. Although this relationship might seem trivial, it can easily be missed in plant tissues, as it requires a local normalisation. To test the second assumption, namely that larger cells are surrounded by smaller cells, we plotted the average of the neighbours’ normalised areas (i.e., the neighbours relative size in regard to its neighbours) against the normalised area of the given cell (Figure 7E). To prevent a circular reasoning (i.e., my neighbour is smaller than me because I am larger than my neighbour), the central cell was always excluded when determining the neighbours’ normalised areas. Nevertheless, we found a very clear relationship supporting the assumption that cells surrounding larger cells are truly smaller than average. This relationship is again much stronger correlated than the normalised area vs area relationship (compare Figure 7C with Figure 7E). Although those two observations together can explain the observed Aboav-Weaire’ s Law-like pattern found in Figure 7A, the underlying mechanism driving the local cell size variation remains unclear, especially since, as pointed out previously, it does not seem to be driven by differences in the number of cell divisions.

Following the effort of linking topology to cell geometry, an important relationship between topology and size in cellular systems was formulated in 1928 by Lewis,

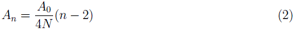

Lewis’ Law empirically relates the number of neighbours with the average area of cells of that topological category; where *n* represents the number of neighbours of a cell, *A*_*n*_ the average area of cells with *n* neighbours, *N* total number of cells, and *A*_0_ the total tissue area. Relating our data in a similar manner, but using normalised area instead, reveals that Equation 2 indeed holds for larger *n* values, but deviates substantially at lower topologies, such as *n*=4 and 3 (Figure 7F). It has been shown that Lewis’ Law can be regarded as the consequence of the equilibrium between entropy and organised form in cellular material, that is, it can be considered to be a direct consequence of the existence of space-filling cells and their topology. Rivier and Lissowski (1982) derived formally, using statistical mechanical considerations, that this law corresponds to the maximal arbitrariness in the distribution of neighbourhood classes of cells that compose a 2D tessellated structure. Their result implies that if a tissue does not follow the relationship Equation 2, such as is the case for out PC tissue, this then indicates that the average area of the cells of this tissue is not simply regulated by the area-filling requirement, but instead other biophysical constraints or biological processes are involved (as also recently shown by Kim et al., 2014). Based merely on these topological and geometrical considerations, we can therefore conclude that the characteristic deviations from both AW-Law and Lewis’ Law that we observe in the experimental data point to the existence of additional regulatory dynamics that are operating during PC development within the leaf. We here propose that they are most likely linked to how divisions are operating in conjunction to mechanics of cell shape establishment.

## Discussion

We revisited theories put forward by D’ Arcy Thompson, armed with novel mathematical and computational approaches and unprecedented potential for data analysis. By analysing a mutant *Arabidopsis* line composed only of pavement cells, we could narrow our focus on the interactions between a similar population of large, non-surface-tension-minimising cells. It allowed us to bypass the natural heterogeneity in patterning and tessellation that is present in a wild type leaf, with it meristemoids and stomata.

Together with advanced microscopy techniques, we could accompany the growth of several leaves, and thereby follow the development of the individuals cells, tracking roughly 50,000 cells and more than 800 cell division events, automatically capturing topological properties amongst the developing cells. We found that, despite the characteristic shapes these cells take up during their development, the topological distribution is conserved between the population of dividing and of differentiating cells. Thus, a topological steady state is reached prior to the PC shape transformation. Despite the pattern being shape- and time-independent, the distribution was unique when compared to other studies: our PC tissue consistently has less 6-sided cells and a higher representation of 5-sided cells than all other animal and plant tissues studied until now (Gibson et al., 2006; Korn and Spalding, 1973; Mombach et al., 1990; Sahlin and Jönsson, 2010).

A large body of theoretical work has focused on investigating the underlying physical and statistical basis that underpin the topological distributions of non-biological and biological materials (Delannay and Le Caër, 1994; Dubertret and Rivier, 1997; Durand, 2015; Durand et al., 2014). Relevant in our context is the result that when it is possible to fix the peak of the topological distribution to *n* = 6 while the extra degrees of freedom allow to vary the variance of the distribution (μ _2_ =< *n*^^2^^ > *-* < *n* > ^^2^^), a useful relationship can be derived between the frequency of 6-sided cells *P* (6) and the variance of the entire frequency distribution μ_2_, by means of assuming a poisson distribution for the possible cellular topologies within the tissue (Le Caë r and Delannay, 1993). The (approximate) relationship that was derived and validated by Le Caër and Delannay (1993) is as follows:

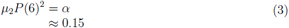

In accordance with this predicted relationship, the experimental data generates α = 0.150049, while the topological model simulation using the Equal division rule without replacement leads to a distribution with α = 0.1558. This further confirms that the relationship between the average frequency of 6-sided cells and the spread of the topological distribution can be captured semi-universally by a single parameter. Further constraining such relationships, Durand et al. (2011) more recently developed analytical models for the statistical mechanics of shuffled two-dimensional cellular tissue to reveal a strict correlation, without any adjustable parameter, between topology and geometry. Their work shows that the standard deviations in the frequency distribution of *n*-sided cells (μ_2_) and in the cell areas themselves (Δ *A*) are in proportion (Durand et al., 2011). In *Drosophila*, this link between area variation and topological variation was experimentally verified and further confirmed through Voronoi models (Sánchez-Gutié rrez et al., 2016). We therefore queried if the pavement cells’ topological distribution could likewise also be linked to the area variation in such a manner. The affirmation that the variation in topology can be correlated unequivocally to the variation in area, however, clearly fails to hold for our system. This follows straightforwardly from the observation that the pavement cell topology frequency distributions are preserved between leaves at different time points (Figure 3A–C), time-points at which the relative variation in cell area varies considerably (Figure S2A–C). This is also true when we analyse the distinct dividing and differentiating cell populations (compare Figure 3D–F to Figure S2D, E). We therefore conclude that for our system a memory-based approach, as suggested by Li et al. (2012), which relies mainly on cell division events, and with non-rigorous constraints on cell surface mechanics, is most appropriate.

Corroborating with this, our topological model, based on graph-simulations, showed that the observed topological distributions could be reproduced by a population of cells that divide with similar rounds of divisions and by dividing in a manner such as to equalise the number of neighbours between the two newly generated daughter cells. Accordingly, on the level of the micro rules which generate these statistical distributions, we were able to find a match between the division events themselves: the experimental data shows a similar trend in the topological redistributions that govern their division planes.

Although we have found a parsimonious model that captures well the macroscopic and microscopic events within the PC tissue, it is not straightforward to interpret these results biologically. A naive and direct interpretation would be to hypothesise that there are mechanisms in place that allow cells to directly assess the number and distribution of neighbours, in such a way that division planes are laid down such as to split the number of different neighbours in two (instead of the contact lengths themselves playing a role). Although there are possible ways, for example through plasmodesmata-mediated cell-cell communication, in which one could envision molecularly defending such a hypothesis, we do not consider it immediately helpful to interpret the results of our topological model in such a manner.

In contrast to the topological model, recently, more attention is being given to the hypothesis that cells assess their shapes and sizes (Chen et al., 1997; Li et al., 2012; Willis et al., 2016) and use such geometric inputs to guide their division planes. What is not so clear, though, is how biological cellular behaviour linked to shape would be able to generate on the topological level the behaviour that we find describes so well the data. Based merely on cell sizes (Figure 7A–F), our data and analysis suggests that purely topological and entropic effects cannot fully explain the resultant topological distributions. For example, the deviations from Lewis’ Law indicate that other mechanisms, of biological origin, are operating. Comparing our PC topological distributions to computer-simulated distributions based on shape-dependent division rules analysed by Sahlin and Jönsson (2010), we found, amongst the very divergent profiles, one clear match as well (Figure S3). In that work, computer simulations based on a vertex-model of equally sized cells with anisotropic growth were performed, in which surface-tension processes act to preserve the areas of the cells and to rearrange the distances between vertexes. Their spatially embedded cellular model prohibits neighbourhood swaps (i.e., T1 events), so it is well-suited to address topological changes through division in plant tissues. Interestingly, contrasting our topological distributions to those generated by their different division hypotheses revealed a large qualitative spread among their results, as well as a large divergence between their experimental *Arabidopsis* WT shoot data and our PC tissue data (Figure S3A). The rule that generated the most closely resembling topological distribution, is the ‘ random split through the Centre-Of-Mass’ rule (Figure S3B). The close correspondence between these distinct implementations of cell divisions suggests that, *under isotropic growth*, surface tension processes coupled with cell growth effectively redistributes cell edges along each individual cell in such a manner that, when divisions occur through the centre of mass in a random direction, this new cell wall tends to split the cell into two equal parts, effectively distributing the neighbours equally between the daughter cells, as performed by our graph model.

However, growth of our *spch* leaf is not isotropic (Figure 7G), nor are the cells of equal size (Figure 7H), consistent with previous careful studies on WT leaf growth dynamics (Donnelly et al., 1999; Kuchen et al., 2012). Thus, it remains an open question what shape-dependent rules – if any – can be mapped onto the topological rules we find here, when taking into account the complex patterns of growth of the *Arabidopsis* leaf. Furthermore, it will depend on future studies that dive deeper into the mechanisms of cell biological control to shed light on which processes are actually being taken into account on the cellular and molecular level that generate the cell division behaviour that we here quantified and which phenomenologically explains the topological distributions. Thus, the quest to link mechanisms between passive biophysics to active cell behaviour based on shape, size and topology — as formalised and initiated by D’ Arcy Thompson a century ago — has still to be finalised, as necessary today as then, in order to unravel how tissues maintain their characteristic and unique properties during growth.

## Materials and Methods

### Confocal images and image processing

*Speechless* mutant (MacAlister et al., 2007) leaves expressing pmCherry-Aquaporin (Nelson et al., 2007) (spch4-pmCherry) were grown and imaged in a custom built perfusion chamber (Calder et al., 2015; Kuchen et al., 2012; Robinson et al., 2011; Sauret-Güeto et al., 2012) from around 7 to 23 days after stratification. Seven different leaves were imaged at in total 121 time-points. Confocal stacks were projected and segmented using custom software and segmented cells were matched between successive time-points. The data is in the form of segmented images, where cell colour corresponds to a unique cell ID, together with tracking data, which links the cell IDs between segmented images. Only cells with a fully defined neighbourhood and not being part of the edge of the leaf (grey cells) were considered in the frequency distributions.

### Cell Division Model

Simulations take the form of operations on a network. An epithelial tissue can be abstracted to an undirected graph, where cells are considered as nodes and links to neighbours are considered as edges. By assuming the tissue has no four-way cell junctions, the number of neighbours a cell has is equivalent to the number of sides it has. With this formalism, all that is needed to completely define a cell division is the cell which is dividing and the two neighbouring cells between which the division plane is placed. The positioning of the division plane depends upon the desired cell division behaviour. For the purposes of this study we considered three different types of behaviour; a randomly oriented division plane, a binomially weighted division plane and an equal split division plane. These three behaviours give different steady state distributions for the final neighbour number frequencies and our study is concerned with which behaviour most closely matches our data. Models were implemented in Python (v2.7) and run on a MacBook Pro (2.2 GHz Intel Core i7, 16 GB RAM).

## Acknowledgements

We thank Matthew Hartley and Tjelvar Olsson for continuous support in the imaging analysis and data processing methods underlying this research. This work has been supported by Consejo Nacional de Ciencia y Tecnología (CONACYT) and by the UK Biological and Biotechnology Research Council (BBSRC) via grant BB/J004553/1 to the John Innes Centre. VAG acknowledges support from the Royal Society Dorothy Hodgkin fellowship.

## Supporting Figures

**Figure S1:**
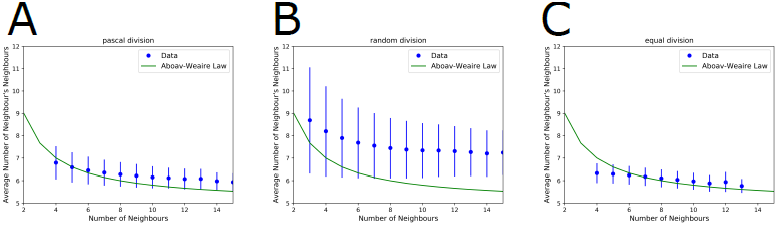
Predicted number of neighbour’ s neighbours in the topology model and Aboav-Weaire’ s Law. Average number of and standard deviation in neighbour’ s neighbours, as a function of neighbour number, for the different topological division rules.

**Figure S2:**
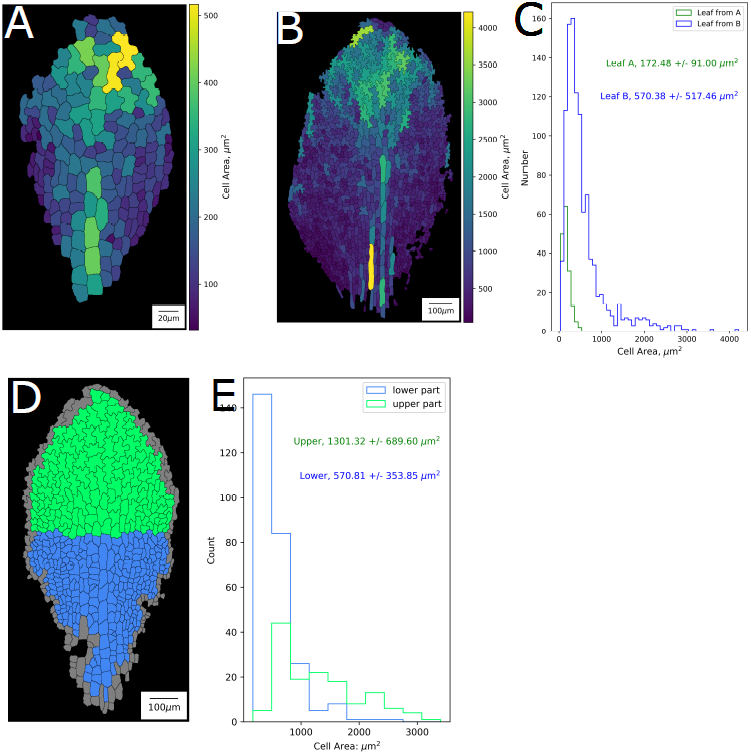
Variation in cellular area over the tissue. (A) Spatial distribution of cellular areas, shown through colour bar to the right of the panel, of a young leaf (time-point 1, 93.25 h after stratification). (B) Spatial distribution of cellular areas of a more mature leaf (time-point 9, 286.50 h). Both tissues correspond to Figure 3A, B. (C) Plot comparing both area distributions, indicating average area and standard deviation. (D) Population of dividing (blue) and differentiating (green) cells, as also shown in Figure 3D.(E) Area distributions of these distinct populations, indicating average area and standard deviation.

**Figure S3:**
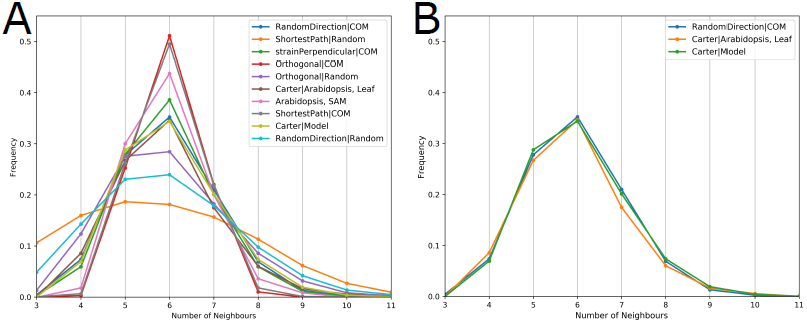
Geometric division-rules compared to PC data. (A) Topological distributions as generated by all the division rules of the vertex-based model presented in Sahlin and Jönsson (2010), compared to the distributions generated by our topological model and observed in our experimental PC data. (B) Only the Random Direction rule bears close similarity to our topological Equal division rule and our PC data.

